# Improving the value of public RNA-seq expression data by phenotype prediction

**DOI:** 10.1101/145656

**Authors:** Shannon E. Ellis, Leonardo Collado-Torres, Jeffrey T. Leek

## Abstract

**Background:** Publicly available genomic data are a valuable resource for studying normal human variation and disease, but these data are often not well labeled or annotated. The lack of phenotype information for public genomic data severely limits their utility for addressing targeted biological questions.

**Results:** We develop an *in silico phenotyping* approach for predicting critical missing annotation directly from genomic measurements using, well-annotated genomic and phenotypic data produced by consortia like TCGA and GTEx as training data. We apply *in silico phenotyping* to a set of 70,000 RNA-seq samples we recently processed on a common pipeline as part of the *recount2* project (https://jhubiostatistics.shinyapps.io/recount/). We use gene expression data to build and evaluate predictors for both biological phenotypes (sex, tissue, sample source) and experimental conditions (sequencing strategy). We demonstrate how these predictions can be used to study cross-sample properties of public genomic data, select genomic projects with specific characteristics, and perform downstream analyses using predicted phenotypes. The methods to perform phenotype prediction are available in the *phenopredict* R package (https://github.com/leekgroup/phenopredict) and the predictions for *recount2* are available from the *recount* R package (https://bioconductor.org/packages/release/bioc/html/recount.html)

**Conclusion:** Having leveraging massive public data sets to generate a well-phenotyped set of expression data for more than 70,000 human samples, expression data is available for use on a scale that was not previously feasible.

## 1 Background

RNA sequencing (RNA-seq) has become the gold standard for assaying gene expression and has advanced our understanding of transcription [1, 2, 3]. To date, tens of thousands of samples have been profiled using RNA-seq technology, and their data have been deposited into the public archive [4]; however, each individual study typically includes only a small number of samples (average study size for human RNA-seq projects using Illumina sequencing within the sequence read archive (SRA) is 24 samples). Without robust sample sizes, reproducibility across studies has been limited [5, 6, 7, 8]. Further, while the scientific community has ensured that these data are publicly available, current repository platforms do not make these data easily accessible for future study. Data in public repositories are often not provided in a consistent format, are not annotated clearly for easy use by independent researchers, or have not all been processed using the same methodology, all of which limits comparability across studies. To assist in improving the utility of the available human expression data in the public repository, we previously developed the `recount2` [9] resource that includes expression data for 70,000 human samples aligned using Rail-RNA [10]. Using a common processing pipeline makes it easier to make direct comparisons across studies. These data, which include expression estimates at the gene, exon, junction, and expressed region levels for each sample, are all available in the `recount2` [9] resource and can be easily accessed using the `R` package `recount`, dramatically reducing previous barriers to accessibility.

While expression information is now available in `recount2` [9], for a large number of these samples, phenotype information is not (Figure 1). For example, while there is expression data available from the sequence read archive (SRA) for 49,657 samples, information for the sex of individual from which the sample was taken is only available for 3,640 (7.3%) samples. Clearly, without such critical sample information, downstream analytic utility is limited. Further, as these data were initially generated in many different labs, analysis using these data must be concerned about unwanted sources of variation affecting their analyses [11]. It is critical to have technical phenotype information available including such as sequencing strategy employee information across samples.

**Figure 1:**
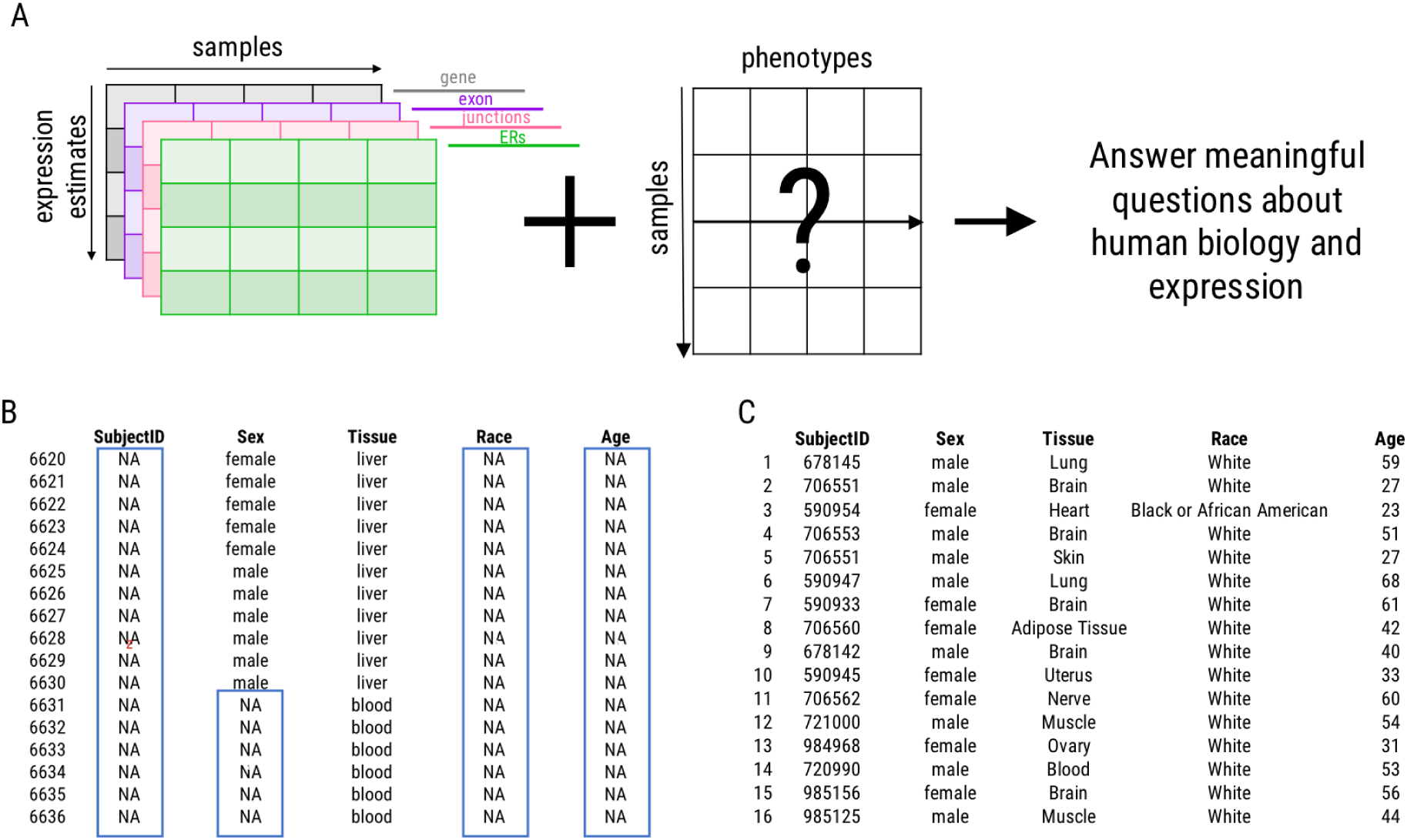
Missing phenotype information. **A.** Phenotype information is critical to answer questions about biology using expression data. **B.** This critical information is missing for many samples within within the SRA (blue boxes). Note that sample phenotype information begins with the 6,620th row, as this is the first row in the dataset for which sex and tissue are available for the same sample. **C.** Missingness is limited within the GTEx data. Expression data from samples with accompanying phenotype information are used to build the predictors. ERs = expressed regions

To address this missing phenotype issue, we have built predictors from the expression estimates themselves for a number of biological and technical phenotypes 2. Using the data within `recount2` [9], which includes samples from the Genotype-Tissue Expression Project (GTEx) [12, 13] (N=9,538), The Cancer Genome Atlas (TCGA) (http://cancergenome.nih.gov/) (N=11,284), and the sequence read archive (SRA) (N=49,657) [4], we have identified regions that are able to accurately predict (1) sex, (2) sample source (whether the sample was generated from a cell line or tissue sample), (3) tissue, and (4) the sequencing strategy (single or paired end) employed. With this critical sample information available for all samples across `recount2`, the utility of the data increases such that accurate downstream analyses (i.e. differential expression analyses) are possible.

**Figure 2:**
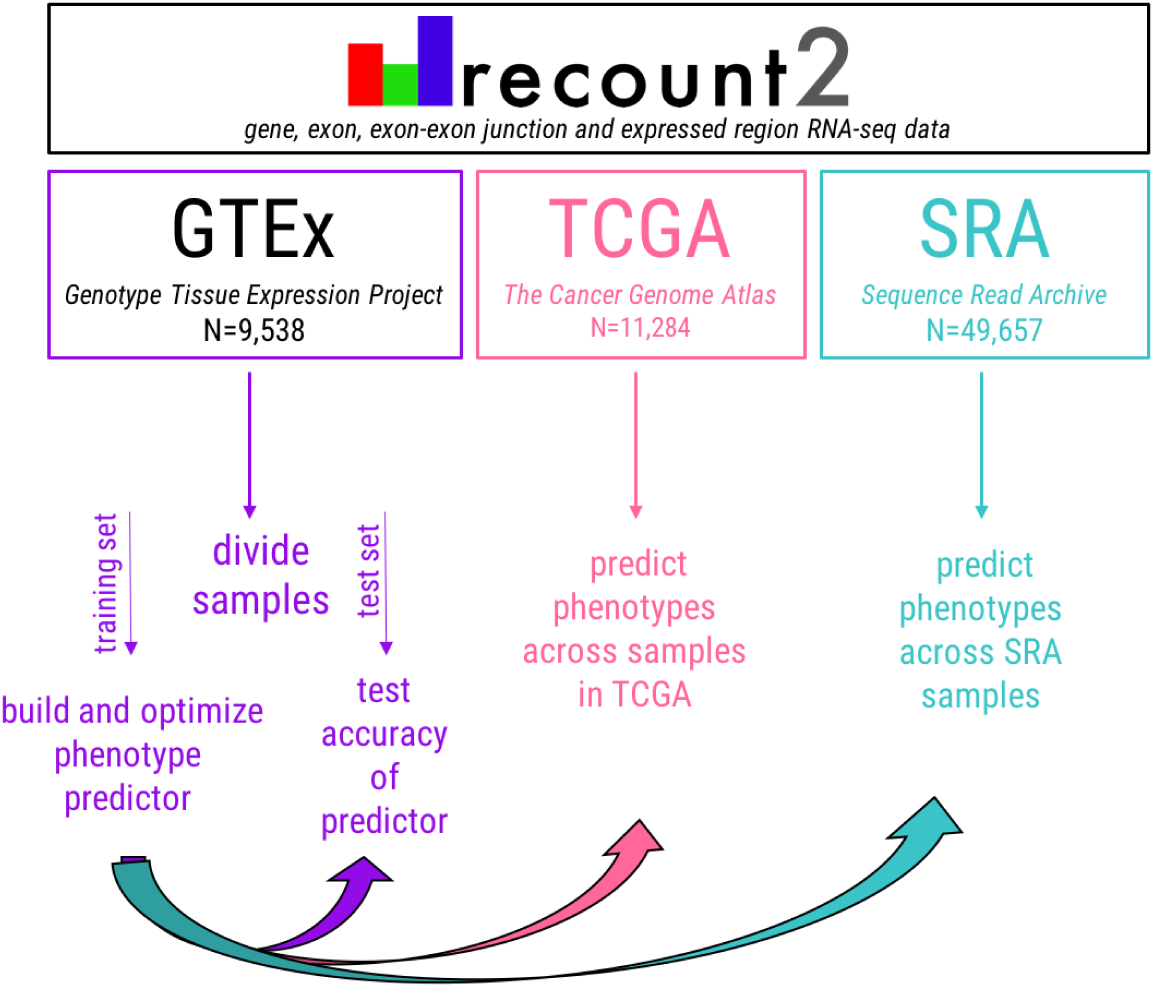
General approach to phenotype prediction. To predict phenotype information, the training data are first randomly divided and the predictor is built. Accuracy is first tested in the training. Upon achieving sufficient accuracy (≥85%), the predictor is tested in the remaining half of the training data set. Phenotypes can then be predicted across all samples within the data set.

Finally, we highlight three possible use cases to demonstrate how the phenotypes predicted can be useful in both their own right as well as for use in future analyses. First, there has historically been a sex bias in biomedical research, such that in both humans and in mice, samples from males are utilized more frequently than females [14, 15]. Accordingly, we demonstrate how predicted phenotypes can be used to characterize the overall breakdown by sex for samples in the SRA. Second, we demonstrate how the phenotypes we have predicted can be used to identify studies whose data may be of particular interest to researchers for their analyses. Last, after identifying a study of interest, we demonstrate how predicted phenotypes can be incorporated into an expression study to gain insight into biology.

## 2 Results

### 2.1 Phenotype prediction

We harnessed the inherent variability of expression data to accurately predict important biological (i.e. sex, sample origin, tissue) and technical (i.e. sequencing strategy) phenotypes from the expression data itself. All phenotype predictors were built using the data available in `recount2` [9]. Data included within this resource include RNA-seq data from (1) the GTEx Project (N=9,538), TCGA (N=11,284), and the SRA (N=49,657). Region-level expression estimates [16, 7], which are annotation-agnostic, were used for prediction. To build the predictors included here and to enable future predictors to be built using this approach, we have developed the R package phenopredict, which includes all the necessary functions to build and test predictors from expression data. Number of both samples used and regions included for predictions included herein are summarized in Supplementary Table S1.

### 2.2 Sex prediction

Sex prediction from RNA-seq data benefits from the biological fact that males and females differ in their sex chromosome composition. To build a predictor for sex from expression data, the GTEx data were first split into a training set (N=4,769) and a test set (N=4,769). The 40 regions (20 each from the X and Y chromosomes) that best discriminated males and females in the GTEx training data were selected and the predictor built. In the GTEx training data, when applying the predictor to the data upon which it was trained, the resubstitution error (see Methods section 4.7) was 0.1%. Sex prediction was then carried out in the GTEx test set, an expression data set in which the data were generated by the same consortia but whose samples were not included to build the predictor. Prediction accuracy was 99.8% in these data. Sex prediction was carried out in data from TCGA (N=11,284), a completely independent and well-characterized data set, where prediction accuracy was 94.2%, and in the SRA samples for which sex information was available (N=3,640), where accuracy was 84.3% (Figure 3A).

**Figure 3:**
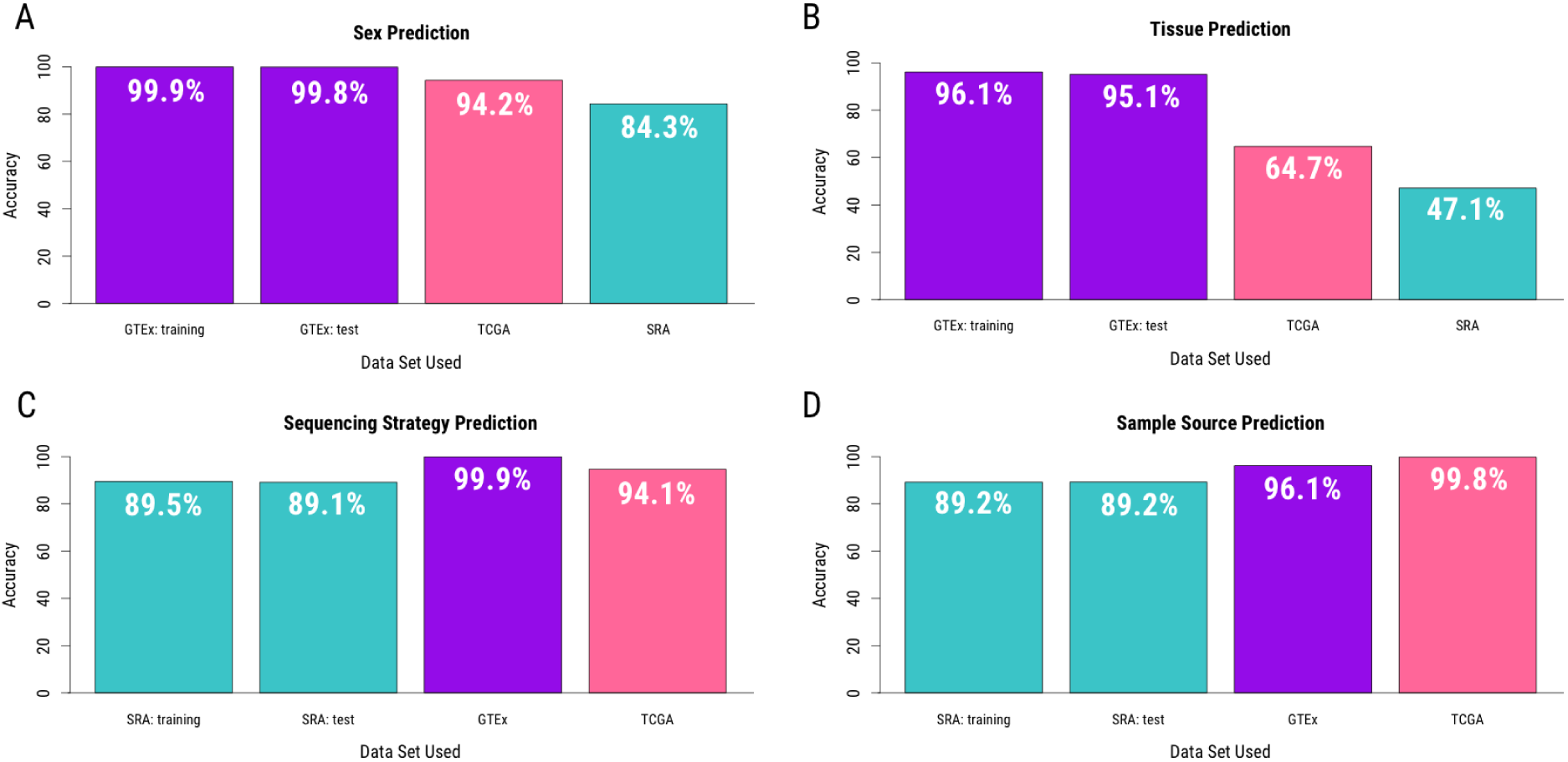
Prediction accuracy. Predictors for critical phenotype information were built from expression data available in `recount2` for **A.** sex **B.** tissue **C.** sequencing strategy and **D.** sample source. Samples for which reported phenotype information is available were used to determine predictor accuracy. GTEx data are in purple, TCGA in pink, and SRA in teal.

### 2.3 Tissue prediction

Tissue prediction is more complex than sex prediction for two reasons: (1) region selection must be carried out across all chromosomes (not just the sex chromosomes) and (2) there are more than two levels for which a predictor must be built. In the case of the data in `recount2` [9], the GTEx training data include information for 30 different tissues. For each tissue, regions with distinct expression profiles relative to all the other tissues in the data set were selected for inclusion in the predictor. For these analyses, 1,057 regions were selected across the autosomes and the sex chromosomes. Accuracy was assessed as above for the GTEx and TCGA data: using the percentage of samples whose tissues correctly predicted from their expression data as our metric. Here, tissue prediction was correct in the GTEx training set 96.1% of the time (3.9% re-substitution error). Accuracy for tissue prediction was 95.1% in the GTEx test set (N=4,769) and 64.7% in the TCGA data (N=7,193) (Figure 3B). We note, however, that TCGA is a cancer data set that includes samples from cancerous tissues as well as healthy control tissue. When this analysis was stratified to separate healthy tissue samples from cancer tissue samples, prediction was accurate 88.6% of the time in healthy tissue (N=613) and 62.5% of the time in cancerous tissue (N=6,705) Supplementary Figure 1), suggesting that altered expression within cancerous tissue samples may benefit from a separate tissue predictor. Assessment in SRA was less direct as tissue information was infrequently available in the metadata available from the SRA. Instead, to find a metric to which we could compare our predicted tissue types in SRA, a natural language processing tool, sharq (http://www.cs.cmu.edu/~ckingsf/sharq/) was employed. sharq was previously used to scrape the abstracts for a subset of studies included in the SRA, extracting its best guess as to the tissue used for study from each. Using this tool, tissue type was assigned to 8,951 SRA samples. These predictions (‘sharq beta’) were previously made available in `recount2` [9]. Comparison between the sharq beta predictions and our tissue predictions from expression data were concordant in 47.1% of the SRA samples (Figure 3B). By using NLP to predict tissue, accuracy within the SRA is likely a lower bound, and one could expect to see improved accuracy, had reported tissue been directly available for comparison.

To assess this claim directly, we compared predictions from the expression data and sharq beta predictions to reported phenotypes for the subset of SRA samples where tissue was reported (N=10,830). Among these samples, there were 336 unique reported tissues within the SRA with 1,442 (13.3%) and 1,496 (13.8%) samples directly matching the predicted tissue from the expression data and sharq beta predictions, respectively. This relatively low concordance often reflects nonuniform or distinct coding within the SRA. Specifically, there are 1,630 cases in which sharq beta and expression predictions are concordant with one another but discordant with what is reported in the SRA. Of these, 1,099 samples (67.4%) include the predicted tissue in the reported tissue (i.e. predicted tissue is ‘brain’; reported tissue is ‘brain mixed human brain’). Of the remaining 531 (531/1,099, 32.6%) samples concordant between sharq beta and expression predictions but discordant with reported tissue, there are a number of explanations. To assess what could be going on, we randomly sampled one sample from each of the 31 unique studies from which these 531 samples were generated. Of these 31 samples, (1) 51.6% of the time (16/31) the predicted tissue were a more general term for the reported tissue (i.e. predicted tissue is brain; reported tissue is ‘dorso-lateral prefrontal cortex’), (2) 25.8% (8/31) of the time the reported tissue was concordant with the predicted tissue, but the reported tissue also included diagnosis (predicted tissue is ‘prostate’; reported tissue is ‘prostate cancer tissue’), (3) 9.7% of the time (3/31) the tissue was coded differently within the SRA for that study (i.e. predicted tissue is ‘blood’, reported tissue is ‘serum’), or (4) 12.9% of the time the predictions are discordant (i.e. predicted tissue is ‘liver’; reported tissue is ‘brain’). Taken together these data suggest that true accuracy is likely between the lower bound of 47.1% determined by comparison to sharq beta predictions and the 82.1% accuracy found in this less rigid comparison to the subset of SRA samples with defined tissue annotations.

### 2.4 Sequencing strategy prediction

Technical sequencing information, such as whether single or paired end sequencing was carried out, is important information for accurate downstream analysis. As the GTEx and TCGA data were all sequenced using a single sequencing strategy, they could not be used to generate the predictor. Instead, to predict this phenotype, the SRA data were randomly split into a training set (N=24,829) and a test set (N=24,828). Eighty regions that distinguished single end sequencing libraries from paired end sequencing libraries were identified and the predictor built. In the training set, the resubstitution error was 10.5%. Accuracy was then assessed in the SRA test samples (N=24,828), GTEx (9,538), and TCGA (N=11,284) samples, where accuracy for sequencing strategy prediction was 89.1%, 99.9%, and 94.1%, respectively (Figure 3C).

While the error rates within the SRA training and test sets are higher than the sex and tissue predictors, this is unsurprising, as this predictor was built in the more heterogeneous SRA samples, where errors in reported phenotypes are more likely and expression data were generated across a number of labs. To test this directly, we compared cases in which predicted and reported sequencing strategy were discordant to samples predicted to have been misreported. Misreported sequencing strategy was calculated independent of expression estimation by comparing the number of reads counted by Rail-RNA [10] and comparing it to the number of spots reported in the SRA. If the sample was reported to be ‘single end’ sequencing but the counted reads value exceeded the number of spots reported, the sample was determined to have a misreported ‘single end’ sequencing strategy when it should have been ‘paired end’. Conversely, for samples labeled paired end where there were exactly twice the number of spots as reads counted, this was determined to be a case where a sample was misreported as ‘paired end’ when it should have been ‘single end’. In the SRA training data, of the 224 samples determined to likely be misreported, 201 (89.7%) were also identified as discordant in the expression predictions, independently confirming misreporting in these studies. Similarly, in the SRA test data, 234 of the 251 (93.2%) samples determined to have been misreported were also discordant between reported and predicted sequencing strategy, further confirming misreporting of sequencing strategy within a subset of the studies in the SRA and suggesting that accuracy reported herein is a lower bound.

### 2.5 Sample source prediction

To utilize data from a human RNA-seq experiment, it is critical the analyst be aware whether the data were generated from tissue directly or from a cell line. To predict this phenotype from the expression data as with the sequencing strategy predictor, regions were selected from the SRA training set. Within this training set, 10,777 samples for which available metadata clearly stated whether the sample came from a cell line (N=4,837; 44.9%) or a primary tissue (N=5,940; 55.1%) were used to select 200 regions upon which the predictor was built. The resubstitution error for this predictor was 10.8%. To assess accuracy in the SRA test set samples, all samples for which sample source information available (N=10,980 including 4,837 cell line and 5,940 tissue samples) were utilized. Prediction accuracy was 89.2% in these data. Applying this predictor across the GTEx (N=9,538) and TCGA (N=11,284) data, demonstrated its accuracy to be 96.1% and 99.8%, respectively (Figure 3D).

### 2.6 Age prediction

In an attempt to predict age, 40 regions associated with age were selected from the GTEx training data using a random forest approach. In the training data, predicted age and actual age were highly correlated (R^2^=0.956); however, in the GTEx training data, these regions were not predictive of age (R^2^=0.101) (Supplementary Figure 2). This initial attempt to predict age was carried out agnostic to sample tissue of origin. To alternatively test if age could be predicted within tissue, regions associated with age were identified across a number of tissues. Given the limited strength of this relationship (Supplementary Figure 3), we did not pursue prediction of age from expression data by this approach further. As such, age predictions have not been included for use. As with age, some phenotypes will likely be better identified through other means – either by manual curation of SRA metadata or through natural language processing [17].

### 2.7 Variance in prediction across data sets

This approach to prediction utilizes simple modeling and interpretable predictors; however, the trade-off for simplicity is that extreme expression estimates in future data sets could pose a problem for accurate prediction. To assess this as a potential issue, we calculated the variance for each of the 40 regions used for sex prediction across each data set as a test case. Two patterns are discernible from the variance across data sets for these 40 regions (Supplementary Figure 4). First, we note that there is generally more variance within the TCGA and SRA data relative to the GTEx samples. As the TCGA samples contain cancer expression data and the SRA samples are extremely heterogeneous, both of which are expected to increase variance in expression estimates, this result is unsurprising. Secondarily, across the 40 sex prediction regions, expression variance is largely similar across data sets at regions used for sex prediction, suggesting that extreme outliers is largely not an issue. However, there are a few regions that demonstrate increased variance in the TCGA and SRA data. Given the overall sex prediction accuracy, this finding suggests that even when more extreme variance is seen within a data set, it does not prohibit prediction. While more extreme variance does not largely affect accuracy within sex prediction, a predictor which directly addresses or controls for these extreme values at the sample level may see improved accuracy.

### 2.8 Assessing discordance

For each predicted phenotype, we report prediction accuracies for samples within each dataset (GTEx, TCGA, SRA). By including information about the accuracy of our predictions we make it possible for downstream models to incorporate this uncertainty. While the overall accuracy for any of the predicted phenotypes assessed was at minimum ≥85%, assessing phenotype accuracy at the sample level is non-trivial. When available, we have included reported phenotype, but this information may not be available for many samples or may be incomplete or incorrect. Phenotype files often use manual data entry and can themselves be prone to error [18]. For example, when we look at the 15 GTEx samples whose reported sex and predicted sex disagree (Supplementary Table S2), 13 of these samples come from the same individual (“GTEX-11ILO”). Further, no samples from this individual had a predicted sex concordant with the sample’s reported sex. Together, this may mean that this sample has mislabeled sex information. The remaining two samples (“SRR809065” and “SRR658331”), each have 12 other tissue samples from the same subject. In both cases, the other 12 samples were predicted to be the reported sex, supporting that these two samples are either incorrect predictions or mislabeled samples that should likely not be used for further analyses.

### 2.9 Use case #1: Sex Analysis within the SRA

There is a long history within the biomedical sciences favoring the study of males relative to females [14, 15]. To assess the prevalence of this bias within the SRA, we have analyzed the overall breakdown of sex in a number of ways. First, we report that overall within the SRA, 58.6% of the samples are predicted to be female, 34.4% are predicted to be male, and 7.4% could not be disambiguated, suggesting that there is no evidence for a male sex bias within the SRA. However, one important caveat is that we do not yet have a handle on how many replicate samples there are within the SRA. Looking specifically at the largest study within the SRA (SRP025982, N=1720) [19], it becomes clear that this study, in assessing the accuracy and reproducibility of RNA-Seq across platforms and labs, analyzed the same samples, which happened to be female, many times. While this is a clear example of a case where replicate samples are biasing the sex breakdown within SRA, we do not yet have an estimate for how widespread this is across all samples within recount2. Having an idea of the number of repeated measures for each individual within the SRA in the future will help to improve understanding of the of overall breakdown sex across the SRA. When broken down by project type, it becomes clear that nearly half of the projects within the SRA (45.3%) include exclusively female samples. Of the remaining projects, 15.7% include exclusively male samples, 1.1% include only samples that could not be disambiguated, and 37.9% of studies include some combination of predicted sexes (Figure 4A). For projects where predicted sex includes samples that are either female and unassigned or male and unassigned, one could reasonably assume that in many of these cases, all samples were likely of one biological sex; however, we will leave it up to future investigators to decide if this is a fair assumption to make for their purposes. Finally, we note, that samples sizes within each study vary from one sample to more than 1500 samples, with the largest study including almost exclusively samples predicted to be female (Figure 4B).

**Figure 4:**
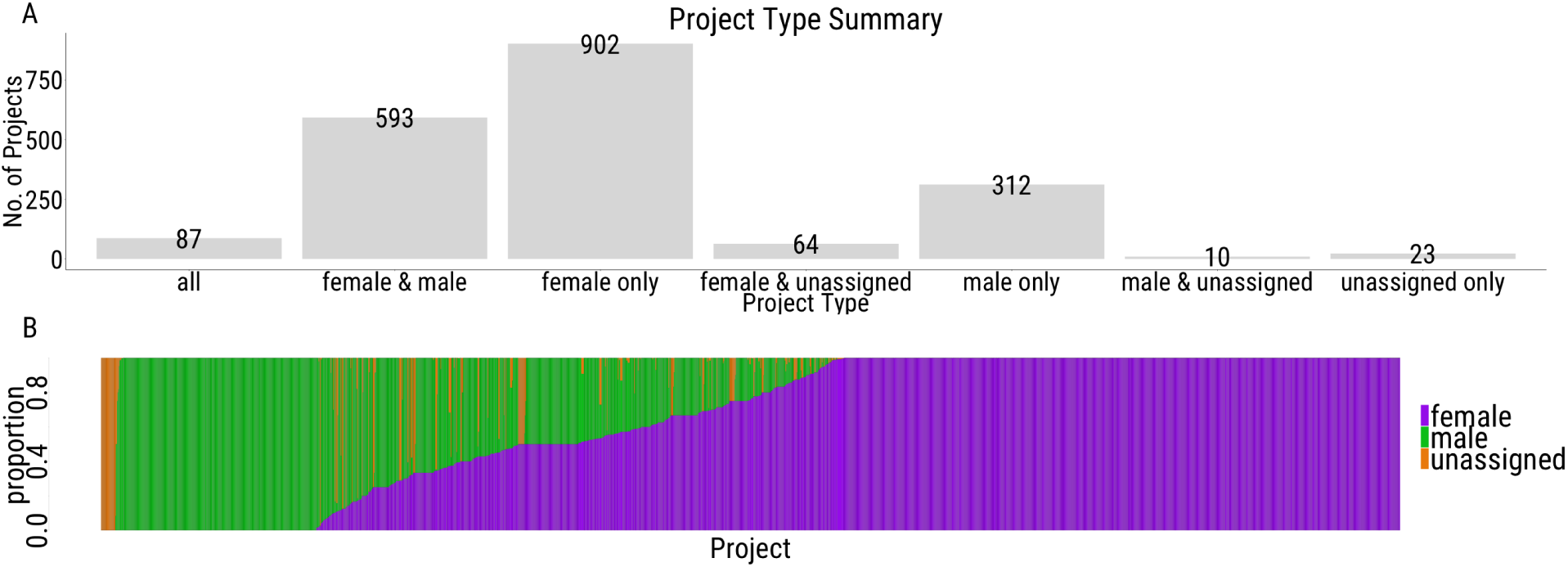
Predicted sex across the SRA. Plots summarize predicted sex across the SRA showing **A.** the number of projects of each project type, and **B.** the proportion of each sex across all projects. Bar width is proportional to study size.

### 2.10 Use case #2: Using predicted phenotypes to identify studies of interest

There is a wealth of expression data within `recount2` [9]; however, with regards to phenotype information, it is only currently possible to filter on information contained within the abstract. Using predicted phenotypes can help to filter out and identify studies of interest to a particular researcher.

For example, we were interested in identifying a cancer study in which sex was not reported. We hoped to assess the effect of including predicted sex in the analysis of these data. To identify such a study, we filtered through cancer studies included in the SRA to identify studies fulfilling the following criteria: expression data (1) predicted to be generated from paired end sequencing, (2) where sex was not reported, (3) where the data were predicted to have come from a tissue (rather than a cell line), and finally (4) that had at least 20 samples. Upon applying these filters, we identified three studies within the SRA meeting these criteria.

### 2.11 Use case #3: Using predicted phenotypes in analyses

Having identified studies of potential interest, we moved forward with study SRP029880 to assess the utility of the phenotypes we predicted (see Methods). This study includes expression data from 54 samples from 18 unique individuals. For each individual, RNA-seq was carried out on a normal colonic epithelial sample (NC), the primary colon cancer epithelium (PC), and a metastatic cancer sampled from the lung (MC). The initial analysis sought to identify gene expression changes that correspond to aggressiveness of colorectal cancer (CRC). To do so, they authors carried out differential gene expression analyses between NC and MC samples (identifying 2,861 significant genes) and between MC and PC samples (identifying 1,846 significant genes). Significance was defined such that *p* < 0.001 and *logFC* ≥ 2. No covariates were included for analysis. The authors then, in an attempt o remove the effects of the fact that the MC samples were sampled from the lung while the PC and NC samples were sampled from the colon, filtered out “liver specific genes (309 genes)” using the TiGER database [20] from their results.

Here, with prediction data available, we can directly and appropriately correct for tissue differences between samples and include sex in the differential gene expression analysis (DGEA). Given the experimental design, we checked to ensure that, for each individual, the same sex was predicted across each of the three samples. Predicted sex was concordant across samples within individual for 17/18 (94.4%) of the samples included for study. (Sample 22 was predicted to be female in one sample and male in the other two). Analysis was first carried out as in the initial publication [21], comparing NC:PC and MC:PC. We note, however, that expression data utilized herein were downloaded from recount2 [9]. Thus, rather than using normalized FPKM, as was done in Kim et al., expression was instead summarized by first scaling the gene counts (see Methods Section 4.14) and then log2 transforming those values. These expression estimates were used to carry out DGEA using the same software and cutoffs as in Kim et al [21]. Despite using the same software, settings, and thresholds used in Kim et al. [21], fewer genes (82.8% and 71.3% less in MC:PC and NC:PC, respectively) were detected to be significantly differentially expressed in our analysis (Figure 5A). As a result, all further comparisons utilized the DGEA results from our analysis, both with and without covariate inclusion to remove the impact of the discrepancy in number of genes identified.

**Figure 5:**
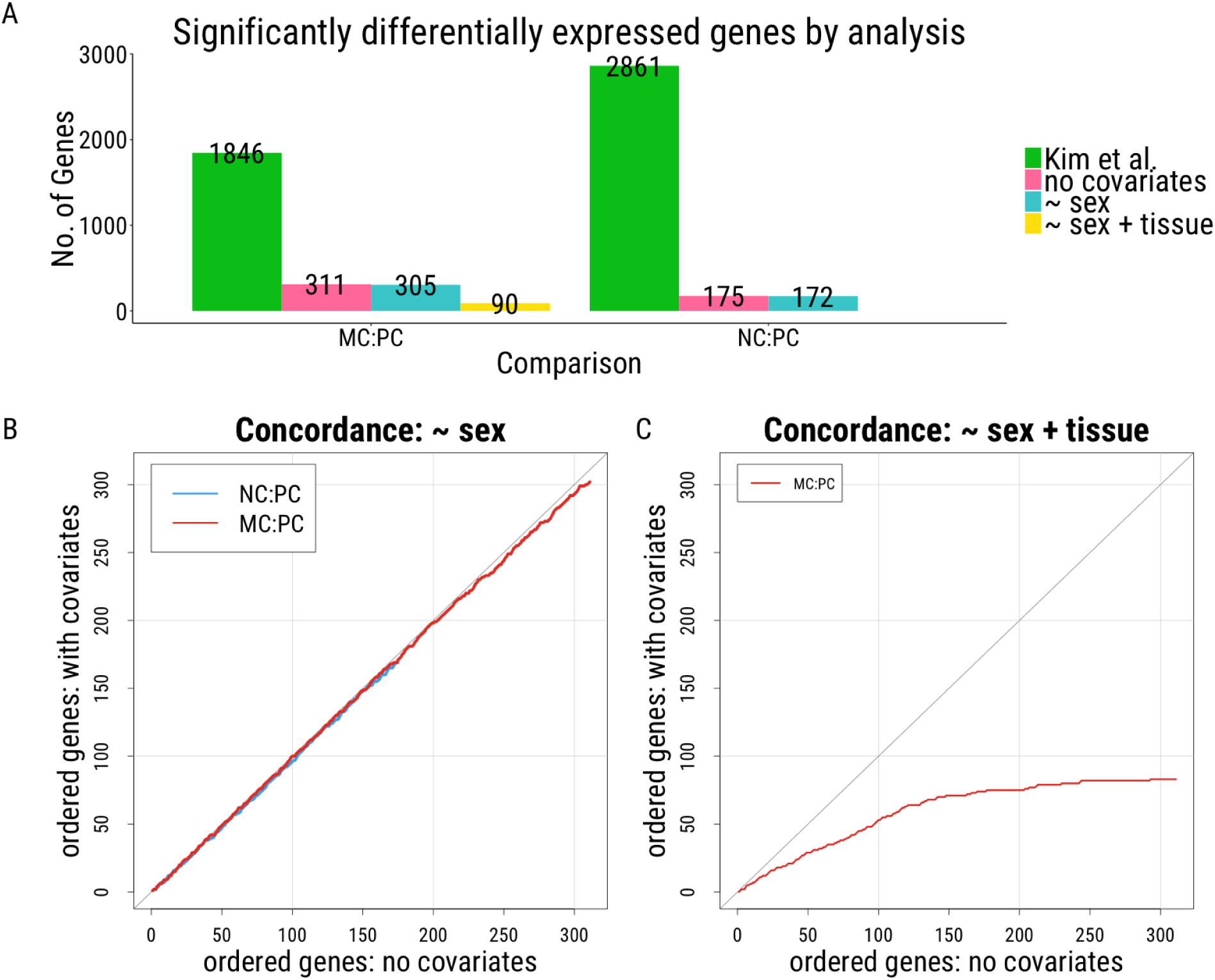
Differential Gene Expression Analysis. **A.** Number of genes reported significant in Kim et al. [21] and the analyses carried out here using their data obtained from `recount2`. **B-C** Concordance at top (CAT) plots [22] comparing DGEA. The number of genes concordant between analyses are plotted, where perfect agreement between analyses’ results would fall along 45-degree line (grey). DGEA where no covariates were included for analysis (x-axis) were compared to **B.** DGEA with sex included as a covariate and **C.** DGEA with both sex and tissue included as covariates. NC = normal colonic tissue; PC = primary colorectal cancer; MC = metastatic cancer (lung)

To assess the effects of covariate inclusion on the analysis, DGEA was carried out with either predicted sex or both predicted sex and predicted tissue as covariates. A final analysis was carried out where both predicted sex and predicted tissue were included as covariates. For each of the NC and PC samples, predicted tissue was either “colon” or “small intestine.” Interestingly, within the MC samples, which were sampled from the liver, seven were predicted to be “small intestine”, the site of the primary lesion, and eleven to be “liver”, the site of the metastases. Interestingly, cancer cell lines do not always recapitulate the tissue from which they were derived, suggesting that cell lines derived from metastases could, in certain cases, recapitulate the primary tissue from which they were derived [23, 24]. Here, these findings suggest, unsurprisingly, that the same may be the case at the tissue level. Thus, if the MC samples were all simply labeled ‘liver’, not only would the results change but the analysis would be completely confounded by tissue. In this case, using predicted phenotypes allows for the most appropriate differential gene expression analysis to be carried out.

The effects of covariate inclusion were assessed using concordance at the top (CAT) plots [22] comparing concordance between genes identified as significant with and without covariate inclusion. Upon inclusion of sex as a covariate in the analysis, the results remain largely unchanged, where 9 of the 10 most significant genes and all 100 of the top 100 genes between analysis are concordant for the MC:PC comparison and 8 of 10 and 96 of 100 genes are concordant for the NC:PC comparison (Figure 5B). However, upon inclusion of both predicted sex and predicted tissue, the results are dramatically different, with concordance at only 6 of the 10 most significant genes and 53 of the top 100 genes between analyses (Figure 5C). As the MC samples were obtained from a completely different tissue, this finding suggests that many (40–47%) of the differences reported as significant between MC and PC samples simply reflect tissue differences between the samples, rather than differences associated with cancer aggressiveness.

## 3 Conclusions

We demonstrate that accurate phenotype prediction from expression data is possible for both biological and technical phenotypes and demonstrate, through three use cases, how predicted phenotypes can be utilized going forward. We have built predictors for sequencing strategy, sex, sample origin, and tissue. These predictors have been applied to the samples within recount2 [9] and can be applied to expression data generated in the future. Predicted phenotypes were generated using the `R` package `phenopredict` and can be accessed in the `R` package `recount` using the `add_predictions()` function. We demonstrate how predicted phenotypes can be used to characterize the samples currently within `recount2`, to identify samples and studies of interest for future study, and how predicted phenotypes can be incorporated into analyses to improve accuracy. Taken together, the availability of this critical phenotype information across the 70,000 samples currently included within `recount2` [9] make downstream analyses with these expression data feasible.

## 4 Methods

### 4.1 Sample summary

Human samples processed using Rail-RNA [10] and incorporated into `recount2` [9] were compiled from the latest (v6) release of Genotype-Tissue Expression Project (GTEx) samples, the sequence read archive (SRA), and the Cancer Genome Atlas (TCGA). The SRA data are comprised of 49,657 publicly available samples from 2,034 distinct RNA-seq projects. The GTEx samples include RNA-seq data for 9,538 samples from 550 individuals and 30 tissues. And, the TCGA samples contributed 11,284 samples from both healthy and cancerous tissue samples. Detailed sample and processing information can be found in Collado-Torres and Nellore et al. [9].

### 4.2 Region level expression data summary

For prediction, as to not limit the sites to expression at annotated regions of the genomes, expressed region (ER) [16, 7] level data were utilized. All GTEx samples were normalized to a library size of 40 million 100 base-pair reads to compute the mean coverage, as available from `recount2` [9]. Expressed regions were defined using a cutoff of 0.5 with the findRegions() derfinder [7] function (https://github.com/nellore/runs/blob/master/gtex/DER_analysis/coverageMatrix/region_bed/find_region_bed.R) resulting in 1,187,643 ERs. The coverage matrix for the ERs was computed using bwtool [25] and custom R code (https://github.com/nellore/runs/blob/master/gtex/DER_analysis/coverageMatrix/bwtool/run_all_bwtool.sh) for GTEx (https://github.com/nellore/runs/blob/master/gtex/DER_analysis/coverageMatrix/bwtool/merge_bwtool.R), SRA (https://github.com/nellore/runs/tree/master/sra/DER_analysis/coverageMatrix/ers_gtex) and TCGA (https://github.com/ShanEllis/covmat_tcga). Expressed region counts were used on the log2 scale. The recount functions expressed regions() and coverage matrix() can be used to reproduce the ERs and coverage matrices. Faster computation of the coverage matrices using `bwtool` can be reproduced using the `recount.bwtool R` package available at https://github.com/LieberInstitute/recount.bwtool.

### 4.3 General approach to prediction

The GTEx data are well-characterized and are comprised of various tissue samples having been generated from the same individual. Given the study design and well-characterized nature of these data, when possible (sex, tissue), the GTEx expression data were used to build the phenotype predictors. When invariability of a phenotype within GTEx limited this approach (sample origin, sequencing approach), the SRA data were used to train the predictor. The first step of our general approach was to randomly sample half of the training data set. Expression data from this subset of individuals comprised our training set. For each predictor, accuracy was then assessed in samples from three tests sets: (1) The remaining samples from the training data set that were not used to build the predictor (2) TCGA, and (3) the data set not used to build the predictor either SRA (in the case of sex and tissue) or GTEx (in the case of sample origin and sequencing strategy). Accuracy was assessed by comparing the predicted phenotype to the reported phenotype in any sample for which this information was available. Upon achieving sufficient prediction accuracy (≥85% in both the training and the first test set defined above), phenotypes were then predicted across all samples within `recount2`.

### 4.4 R package: phenopredict

We have built the R package phenopredict which can use expression data and phenotype information from any study to build a phenotype predictor. Functionality within this package allows for (1) regions to be selected for the phenotype of interest (`select_regions()`), (2) a predictor to be built for either continuous or categorical variables (`build_predictor()`), (3) the predictor in to be tested on the training data to assess resubstitution error (`test_predictor()`), (4) for data to be extracted from a new data set for the same regions upon with the predictor was built (`extract_data()`), and (5) for phenotype prediction in this new data set (`predict_pheno()`). Additionally, for analyses in which data have to be extracted across multiple files (i.e. across chromosomes), there is a function (`merge_input()`) to merge across the output of `select_regions()` prior to building the predictor.

### 4.5 Region selection: select_regions()

Within the training data, where phenotype information is available, the first step is to identify expressed regions associated with the phenotype of interest. To accomplish this, `select_regions()` in `phenopredict` was used.

**Continuous variables:** Regions are identified within `select_regions()` by employing a linear model within `limma’s lmFit()` framework [26]. In this framework, phenotype is regressed upon expression such that:

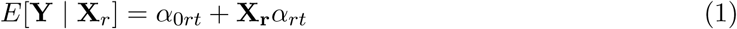

where **Y** is the continuous phenotype. **X**_*r*_ is the design matrix with the expression values for each region and optionally other covariates, and *αo_tr_* and *α_rt_* are the coefficients relating the expression of region *r* to phenotype *t.* To ensure predictors herein were as generalizable across data sets as possible, no covariates were included during region selection; however, functionality exists within `phenopredict` to include covariates. We use `limma` to moderate variances across the regions tested, thus shrinking region-wise sample variances. These moderated variances are utilized in identification of regions most highly associated with the phenotype of interest. We then pick the top *s_t_* most significant regions using the `topTable()` function from `limma`. The number of regions to select is specified by the user.

**Categorical variables:** The same approach as above (equation 1) is carried out for each level in the phenotype. For each level of the phenotype we create a vector of indicator variables ***Y***_*t*_ such that *Y*_*it*_ is an indicator that is one if sample *i* is level *t* and zero otherwise. We then follow the modeling procedure described above to select the top regions.

### 4.6 Building the predictor: build_predictor()

**Categorical variables:** Coefficient estimates are calculated for each level in the phenotype of interest. Using expression from the selected regions and the phenotype of interest as input, `validationCellType()` from the `R` package `minfi` [27] is used to fit the model:

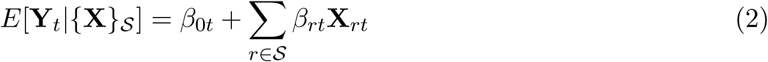

where *S* is the set of all selected regions across levels of the phenotype for categorical variables. We fit the model subject to the condition that *β_rt_ >* 0∀*β*_*rt*_.

**Continuous variables:** A parallel random forest is used to build the predictor in the training data where the features are the selected expressed regions **{X}_s_** and the outcome is the phenotype vector **Y**.

Random forests for continuous variables were constructed using `parRF` method within the R package `caret` [28]. The model was provided with both expression estimates for the selected regions and covariate information, if desired. In the case of age, prediction was attempted both with and without covariate (sex, BMI, and tissue) inclusion, to ensure accuracy could not be improved upon covariate inclusion. The fits from this model are output for prediction.

### 4.7 resubstitution error: test_predictor()

**Categorical variables:** To assess prediction accuracy in the training data, where truth is known, predictions are made from the predictor built in equation 2. `test_predictor()` makes predictions from the expression data and coefficients generated in `build_predictor()` using the framework established in `projectCellType()` from minf [27, 29]. For each sample, the likelihood the sample is each level in the phenotype is calculated. Predicted phenotype is subsequently assigned to the most likely level within the phenotype. If two levels within the phenotype receive the same estimate, the sample is predicted as “unassigned.” Finally, this function reports (1) predicted phenotypes, (2) reported phenotypes, and (3) resubstitution error, where:

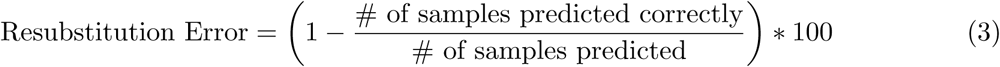

**Continuous variables:** Predictions are made from the random forest model’s fits. Here, this was carried out using `predict()` in `R.` Accuracy was assessed using the correlation coefficient (*R*^2^) and root mean square error (RMSE) between the reported and predicted values.

### 4.8 Extracting regions in a new data set: extract data()

Before prediction can be carried out in a new data set, expression data at the precise regions used to build the predictor must be extracted. `extract_data()` can be used to obtain expression estimates in a new data set at the same eact regions selected by `select_regions()` and used to build the predictor. By using expression from a fixed set of genomic regions in the new data set, rather than identifying a new set of predictive regions, we limit the possibility of overfitting our model.

### 4.9 Phenotype prediction: predict_pheno()

**Categorical variables:** Phenotype prediction in a new data set occurs as in `test_prediction()` with the exception that expression data in this case comes from a new expression data set (*X_new_*). These expression data are from an independent set of samples but include information for the same regions (*r*) as were characterized in the data set (*X_r_*) used to build the predictor.

**Continuous variables:** Fits calculated by `build_predictor()` from the random forest model are then applied to these data and predicted phenotypes are provided.

### 4.10 Assessing accuracy

Categorical variables: The subset of samples within each data set for which reported phenotypes were available were used to assess accuracy, such that:

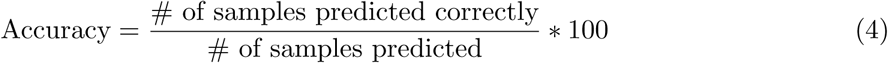

Here, the number of samples predicted includes all samples for which phenotype information was available and whose reported value was included in our predictor (see Supplementary Table S1). Using sex as an example, all SRA samples whose sex was reported as “male”, “M”, “Male”, “m”, “female”, “F”, “Female”, or “f” were included in the denominator. However, samples whose reported sex was missing (NA) or whose reported value was not meaningful (i.e.“not collected”, “not determined”, “Unknown”, etc.) were not included in the calculation of accuracy. Sex was, however, predicted across all samples after accuracy assessment.

**Continuous variables:** Accuracy was again assessed using the correlation coefficient (*R*^2^) and RMSE between the reported and predicted values, for samples where information was available.

Ellis et al.

### 4.11 Integration with recount2

Predicted phenotypes are available through the R package `recount`. Predicted phenotypes can be included using the the `add_predictions()` function. For each predicted phenotype, this function adds reported phenotype, predicted phenotype, and prediction accuracy. The most recent release will be used by default; however, older versions will remain accessible through recount. An example of this function is provided at https://gist.github.com/ShanEllis/?add_pred.R and demonstrates how to add predicted phenotypes to the GTEx data within `recount`.

### 4.12 Use case #1: Sex analysis within the SRA

Sex within the SRA was studied across the 49,657 samples in `recount2`. Predicted sex was first summarized overall, where each sample received a predicted sex of male, female, or unassigned (when sex could not be disambiguated). Secondarily, sex was summarized by project. When all the samples within a project were of the same predicted sex, they were classified accordingly as female only, male only, or unassigned only; however, projects with samples of more than one predicted sex were determined to be ‘mixed’. Of these mixed studies, sex was summarized within each possible combination: all (male+female+unassigned), female+male, female+unassigned, and male+unassigned. Finally, sample size broken down by sex within each project type was summarized.

### 4.13 Use case #2: Using predicted phenotypes to identify studies of interest

Predicted phenotypes can be used to identify samples or studies of interest for future study. To exemplify this, constraints were placed on the predicted phenotype data to search for a study within the cancer studies in `recount2` requiring that possible projects must (1) employ paired RNA-seq, (2) not have reported sex included in the study’s metadata, (3) have tissue a predicted sample source (rather than be derived from a cell line), and (4) have at least 20 samples in the study. Studies fitting these criteria were considered for use in Use case #3.

### 4.14 Use case #3: Using predicted phenotypes in analyses

After identifying two studies in case study #2 that fit our criteria, the corresponding abstracts for each study were searched within `recount2` [9]. Not only did SRP027530 have a smaller sample size (*N_SRP027530_* = 20; *N_SRP029880_* = 54), but the abstract of these data included the breakdown of females and male samples [30]. Thus, despite the fact that sex was not directly reported in and accessible from the SRA metadata, a quick search made it clear that sex could be relatively easily obtained for this study. As we were interested in studying the effects of sex on an analysis that otherwise did not include sex in their analysis, we moved to study SRP029880.

Gene-level count data were downloaded from `recount2` [9] for project SRP029880. Raw counts were scaled to the total coverage of the sample. Cancer status (NC, PC, MC) and sample id were obtained from the reported SRA metadata field ‘title’. Differential gene expression analysis was first carried out as reported in the initial publication [21]. Briefly here, DGEA was carried out using the edgeR [31] R package, which assumes a negative binomial distribution for expression to detect differentially expressed genes. Both normal colon epithelium (NC) and lung metastases (MC) were compared to primary colorectal tissue (PC) samples. To recapitulate the analysis from the initial publication, no outlier genes or samples were removed, no covariates were included, and no adjustment was made for the fact that the PC samples were used in both comparisons. As was done in the initial publication, to protect against hypervariable genes driving differences detected gene dispersion was estimated for the log2 scaled gene counts for 58,037 genes across 54 samples (18 unique individuals). For each gene (using the estimated dispersions calculated), and each comparison (NC:PC, MC:PC) a negative binomial generalized log-linear model (glm) was fit. To be in line with the initial publication [21], genes were considered statistically significant if the p-value was <0.0001 and the log fold change (logFC) was ≥2.

To assess the effects of covariate inclusion on this analysis the same analysis as above was carried out with the exception that covariates were included. First, the inclusion of predicted sex as a covariate on the analysis was assessed. Secondarily, the inclusion of predicted sex and tissue was assessed. In line with expectation, within this study, samples were predicted to be either ‘small intestine’, ‘colon’ or ‘liver’. Samples predicted to be either ‘colon’ or ‘small intestine’ were combined into the umbrella category ‘intestine’. DGEA was carried out as above. To note, however, tissue only varies in the MC:PC comparison, where both liver and colon samples were analyzed. Accordingly, this analysis was carried out for the MC:PC comparison and not NC:PC, where all samples were from the colon.

To compare across the three DGEA (no covariates, sex, and sex + tissue), for each of the two cancer comparisons (NC:PC and MC:PC), concordance among genes determined to be significantly differentially expressed were assessed. Concordance at the top (CAT) plots [22], where the number of significant genes in common between the analyses being compared is visualized, were generated.

### 4.15 Availability of data and materials

Analyses were all carried out in `R 3.3.1.` Code for the `R` package `phenopredict` is available at https://github.com/leekgroup/phenopredict. Code for prediction of each phenotype is available at https://github.com/ShanEllis/phenopredict_phenotypes. Use case code is available at https://github.com/ShanEllis/phenopredict_usecase.

## 4.16 Acknowledgements

We would like to thank Andrew Jaffe, Kasper Hansen, and Abhinav Nellore for helpful discussions. SEE, JTL, and LCT were partially supported by NIH R01 GM105705.

## 4.17 List of Abbreviations

RNA-Sequencing (RNA-seq); Genotype Tissue Expression Project (GTEx); The Cancer Genome Atlas (TCGA); Sequence Read Archive (SRA); expressed region (ER); generalized log-linear model (glm); log fold change (logFC); colorectal cancer (CRC); normal colon epithelium (NC); lung metastases (MC); primary colorectal tissue (PC); Differential gene expression analysis (DGEA); Root mean square error (RMSE); concordance at the top (CAT)

## Supplemental Materials

**Figure 1:**
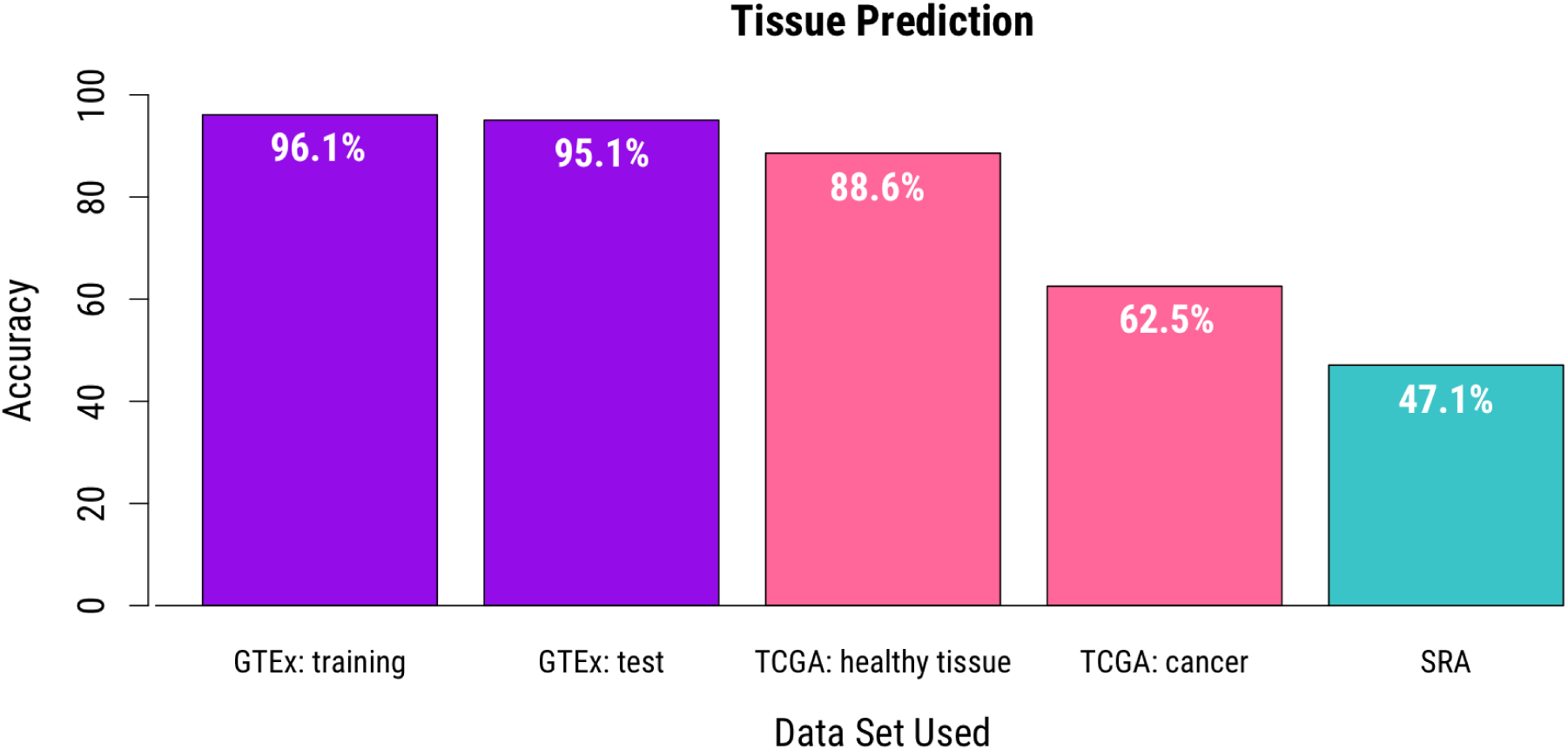
Tissue prediction within TCGA. Prediction accuracy within TCGA (pink) shows marked improvement within healthy tissue relative to cancer tissue samples. GTEx data are in purple, TCGA in pink, and SRA in teal.

**Figure 2:**
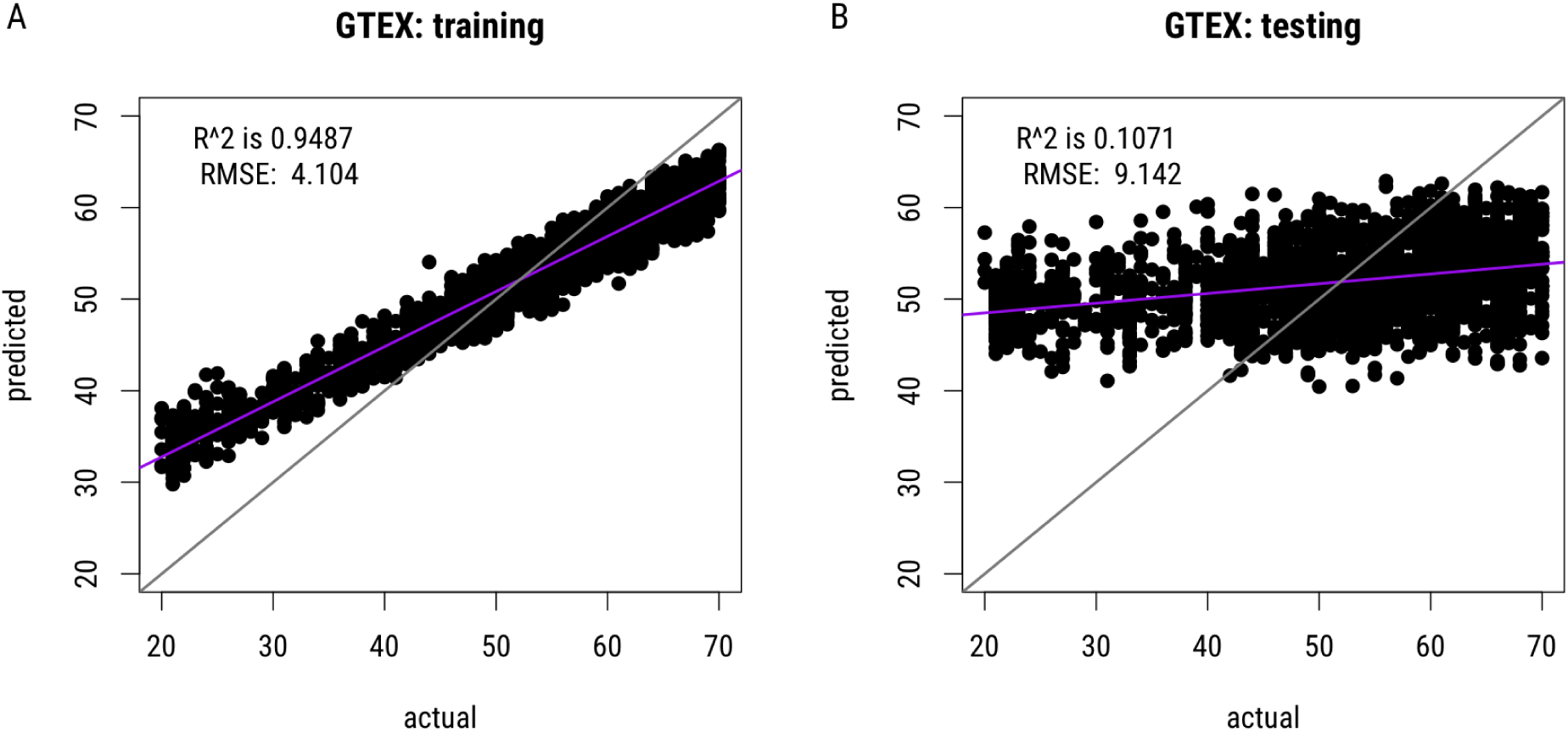
Age Prediction. Parallel random forest was used to build a predictor for age from expression data. Predicted ages vs. actual (reported) age (in years) are shown in the A. training and B. test data sets. Correlation (R^2^) and root mean squared error (RMSE) are shown for each data set.

**Figure 3:**
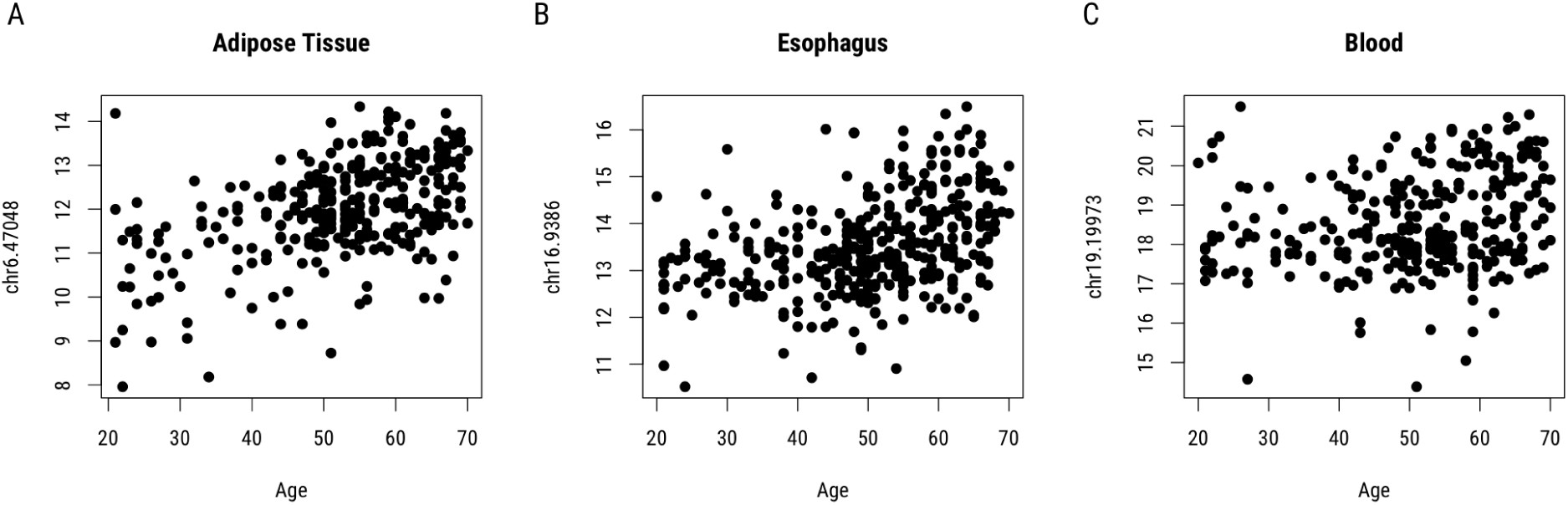
Expression within tissue. Expression estimates (log2 scale) for the region most highly associated with age (years) in A. adipose tissue B. esophagus and C. blood are shown.

**Figure 4:**
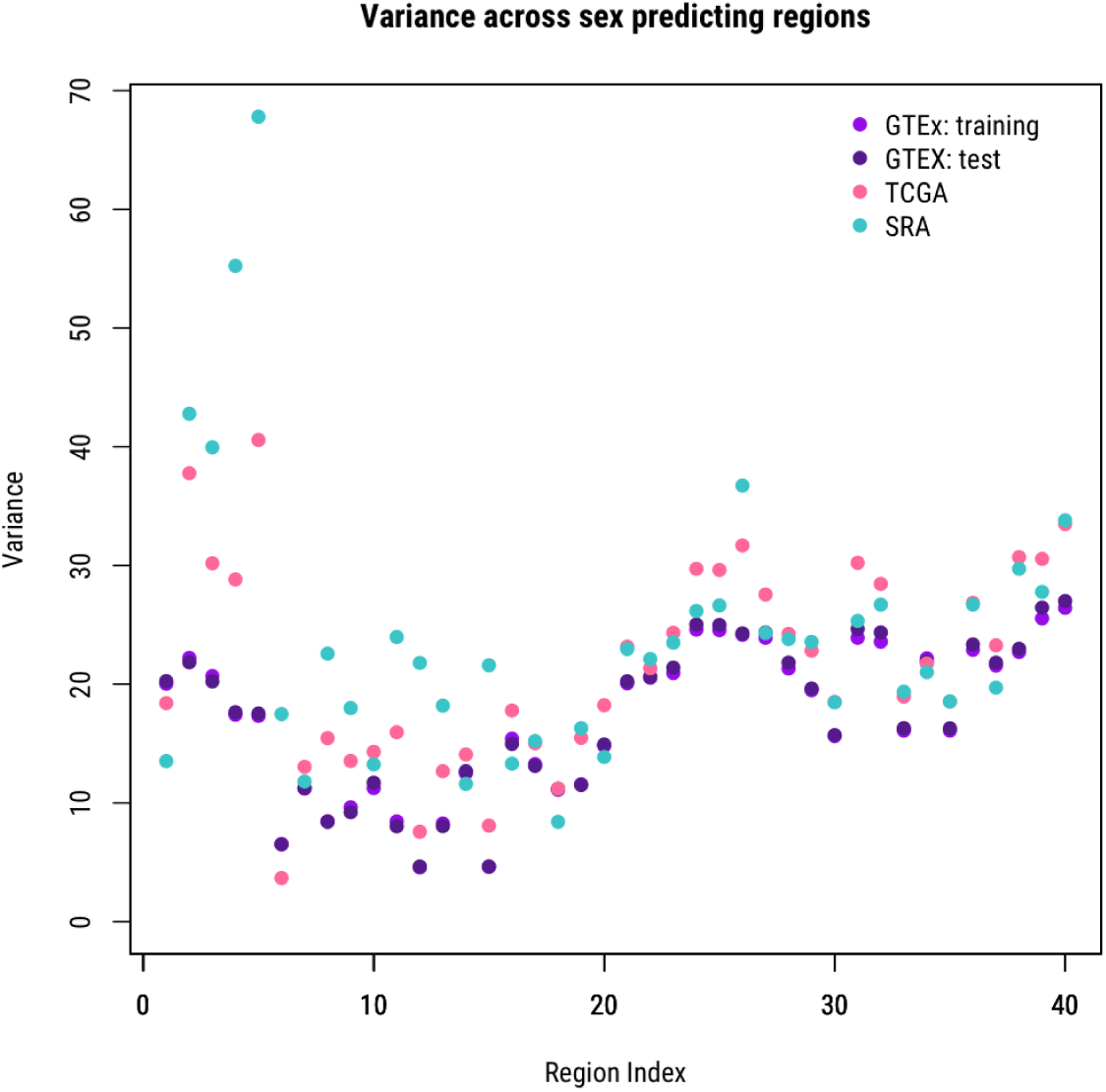
Variance across sex prediction regions. Variance in log2 expression at the 40 regions used to predict sex are shown across the data sets within `recount2`. GTEx training and test sets are in purple, TCGA in pink, and SRA in teal. Indices 1–20 correspond to regions on the X chromosome, while 21–40 correspond to the Y chromosome.

**Table S1:** Phenotype predictor summary

| phenotype | # regions | training set | N<sub>training</sub> | test set #1 | N<sub>test</sub> #1 | test set #2 | N<sub>test</sub> #2 | test set #3 | N<sub>test</sub> #3 |
| --- | --- | --- | --- | --- | --- | --- | --- | --- | --- |
| sex | 40 | GTEX: training | 4,769 | GTEX: test | 4,769 | TCGA | 11,245 | SRA | 3,640 |
| tissue | 1,057 | GTEX: training | 4,769 | GTEX: test | 4,769 | TCGA | 7,193 | SRA | 8,951 |
| sequencing strategy | 80 | SRA: training | 24,829 | SRA: test | 24,828 | TCGA | 9,929 | GTEX | 9,538 |
| sample source | 200 | SRA: training | 10,777 | SRA: test | 10,980 | TCGA | 9,929 | GTEX | 9,538 |

**Table S2:**
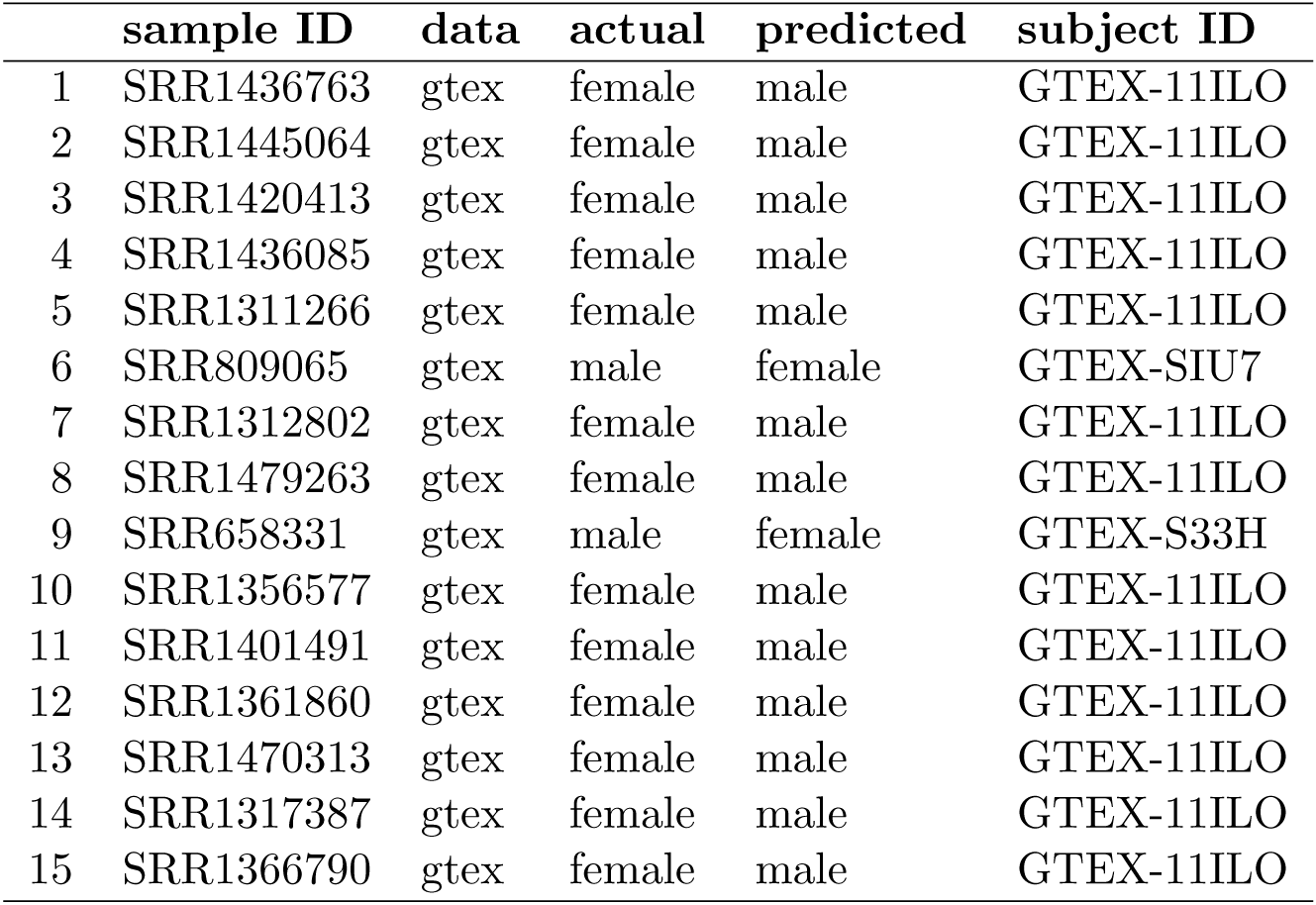
Discordant sex prediction within GTEx

